# Targeting a malaria merozoite surface protein with mRNA vaccine generates multifunctional antibodies

**DOI:** 10.64898/2026.03.26.714647

**Authors:** Adam Thomas, Tom Rünz, Timothy Ho, Stewart Fabb, Chee Leng Lee, Sandra Chishimba, Rekha Shandre Mugan, Linda Reiling, Liriye Kurtovic, Claretta D’Souza, Colin Pouton, James Beeson

**Affiliations:** Burnet Institute, Victoria, Australia; Department of Immunology, Monash University, Victoria, Australia; Faculty of Mathematics and Natural Sciences, University of Bonn, Germany; Department of Microbiology, Monash University, Victoria, Australia; Monash Institute of Pharmaceutical Sciences, Victoria, Australia; Department of Medicine, University of Melbourne, Victoria, Australia; Department of Medicine and Infectious Diseases, University of Melbourne, Victoria, Australia

## Abstract

**Introduction:** Malaria is a leading health problem with high disease burden and mortality rates worldwide. Currently approved vaccines target the sporozoite form of *Plasmodium falciparum* that initially infects the liver, but only provide modest protection against malaria in young children. There is an urgent need to develop next-generation malaria vaccines that target multiple parasite developmental stages for greater efficacy. Antibodies to merozoites, which are involved in blood-stage replication, and are associated with clinical illness, have multiple functional activities and can protect against malaria. A promising merozoite vaccine candidate is Merozoite Surface Protein 2 (*Pf*MSP2). Antibodies to *Pf*MSP2 can promote multiple antibody Fc-mediated functional activities to clear merozoites.

**Methods:** We developed and evaluated monovalent and bivalent (3D7 and FC27 variants) *Pf*MSP2-based mRNA vaccines. We designed and codon-optimised mRNA, which was validated for *in vitro* expression in mammalian cells, and subsequently formulated as lipid nanoparticles for vaccination of mice in a 3-dose regimen. Vaccination with recombinant *Pf*MSP2 protein with adjuvant was performed for comparison. We evaluated the induction of antibodies and functional activities relevant to protective immunity.

**Results:** mRNA vaccines induced prominent IgG responses using monovalent (3D7 allele) and bivalent (3D7 and FC27 alleles) vaccines encoding near full-length *Pf*MSP2, and antibodies recognised the surface of whole merozoites. Vaccine responses were equivalent to, or superior than, a recombinant protein-based *Pf*MSP2 vaccine. The bivalent vaccine induced equivalent antibodies to the two *Pf*MSP2 alleles. Vaccination induced cytophilic IgG subclasses with multiple functional activities, including complement fixation, binding of human Fcγ-receptors I and IIa, and opsonic phagocytosis.

**Conclusions:** *Pf*MSP2 is highly immunogenic using the mRNA vaccine platform and induces antibodies with multiple functional activities associated with protective immunity in humans. Combining *Pf*MSP2 with other merozoite and sporozoite antigens is a promising strategy to develop highly efficacious vaccines to achieve malaria control and elimination goals.

## Introduction

Malaria has a high global health burden spread across 83 countries and is transmitted by *Anopheles* mosquitoes, with the highest morbidity occurring in Africa. *Plasmodium falciparum* is the parasite species responsible for most deaths globally [1]. The World Health Organisation (WHO) estimated more than 280 million recorded cases and nearly 610,000 deaths in 2024, with the majority being children under 5 years. Progress in reducing malaria has stalled since 2015 [1], in part due to increasing resistance to antimalarial drugs and insecticides [2, 3], with risks that the burden will increase in coming years. Therefore, effective vaccines are required to complement current interventions to reduce the malaria burden and achieve and sustain malaria elimination in the long-term. Next-generation vaccines are required to achieve the Global Technical Strategy (GTS) targets of reduction in global malaria incidence and death rate of 90% by 2030 compared to the 2015 baseline [1].

The *P. falciparum* lifecycle is complex, whereby the sporozoite parasite form is transmitted to the skin from an infectious mosquito bite. The sporozoites establish infection in the liver and develop into merozoites, which are released into the blood and undergo asexual replication within red blood cells. This results in an exponential increase in parasitemia and clinical malaria symptoms. The only approved vaccines for malaria, RTS,S/AS01 and R21/MatrixM, target the exact same sporozoite antigen and aim to prevent hepatic infection [4]. However, several challenges remain for these vaccines: efficacy is modest and variable, immunity wanes relatively quickly and annual boosters are needed, efficacy varies by population and is impacted by *P. falciparum* strain diversity, and efficacy is substantially lower in older children and adults [5–8]. Next-generation vaccines targeting both the sporozoite and merozoite could be superior in reducing malaria morbidity and mortality rates and achieve greater efficacy. Merozoite antigens are attractive targets for vaccine development as they have the potential to reduce or prevent blood-stage replication of *Plasmodium* species thereby reducing or preventing clinical illness. Antibodies to merozoites can function by directly inhibiting merozoite invasion of erythrocytes and through antibody Fc-mediated functions including complement fixation and activation and interactions with Fc-receptors to promote opsonic phagocytosis and cellular cytotoxicity with monocytes, neutrophils and natural killer cells [9–13]. Recently, a leading merozoite vaccine based on the *Pf*Rh5 antigen has shown modest efficacy (50% over 6 months) against malaria in young children [14], and functions by inhibition of merozoite invasion. Efficacy could potentially be improved by generating antibodies with multiple effector functions [13, 15, 16]. Combining *Pf*Rh5 with another merozoite antigen that can generate antibodies to the merozoite surface to promote complement fixation, opsonic phagocytosis and cellular cytotoxicity is a potential strategy to achieve higher protective immunity.

*P. falciparum* Merozoite Surface Protein 2 (*Pf*MSP2) is a promising candidate for blood-stage vaccine development [12, 17, 18]. *Pf*MSP2 is a highly abundant surface protein on the merozoite that is tethered to0020the membrane by a glycosyphophatidylinositol (GPI) anchor [19]. Naturally acquired antibodies to *Pf*MSP2 have been associated with protection against malaria across different populations [20]. These antibodies can function via the antibody Fc region to mediate complement fixation and opsonic phagocytosis leading to inhibition of blood-stage replication and clearance [11, 12]. Complement fixation by antibodies against merozoites can inhibit invasion and lead to merozoite killing [11, 12], and complement fixing antibodies to *Pf*MSP2 have been associated with protection [21]. Antibodies to *Pf*MSP2 do not directly inhibit blood-stage replication *in vitro* on their own, but can fix and activate complement to inhibit merozoite invasion and parasite replication [11]. A phase 2b trial of a *Pf*MSP2-based vaccine in a malaria-endemic population showed evidence of strain-specific vaccine efficacy. *Pf*MSP2 has a large variable region (∼220 residues) flanked by conserved N- and C-terminal domains [22]. These variable regions contain repetitive dimorphic sequences that give rise to two allelic families (3D7 and FC27) [22]. Therefore, diversity in *Pf*MSP2 can be covered by inclusion of the two major alleles in a vaccine to account for high sequence diversity and polymorphisms. A phase 1 clinical trial in malaria-naïve adults with a bivalent adjuvanted recombinant *Pf*MSP2 vaccine containing both alleles showed significant induction of multifunctional IgG that targeted all *P. falciparum* strains [11, 12, 17].

Advances in molecular technologies offer new opportunities for next-generation vaccine development such as SARS-CoV2 mRNA-LNP vaccines manufactured by Moderna and Pfizer that have been widely used globally [23]. The mRNA platform has the advantage of efficient multi-antigen formulations and adaptability to emerging pathogens and strains [24]. The mRNA-LNP vaccine formulations induce robust antibody responses with small doses for sufficient immunogenicity to various infectious diseases [24–27]. Here, we designed and evaluated *Pf*MSP2 vaccines using the mRNA-LNP platform with the aim of generating multifunctional antibodies against merozoites. This work provides a proof-of-concept for including merozoite antigens that generate Fc-mediated functional immunity against merozoites in vaccine design, and the potential for including *Pf*MSP2 in multi-antigen mRNA vaccines for malaria.

## Results

### PfMSP2 mRNA vaccine shows high immunogenicity in mice

We first designed an mRNA vaccine construct based on the 3D7 allele of *Pf*MSP2, which was codon-optimised and truncated (Δ19N and Δ23C) to remove the N-terminal secretory signal and C-terminal GPI anchor sites, respectively (Figure 1A). This design was identical to the recombinant *Pf*MSP2 used in a previous human clinical trial, and was used in this study as a control regimen [17]. To confirm protein expression, we transfected Expi293F (HEK293) suspension cells with the naked mRNA and samples were collected and processed under reducing conditions for western blot analysis. The *Pf*MSP2 3D7 mRNA showed increasing protein expression over time that was detected using antigen-specific antibodies (Figure 1B). We then formulated our construct as lipid nanoparticles (LNPs) for preclinical testing in female C57BL/6 mice (n=5 per group) which received a fixed-dose (3 × 5 μg) or dose-escalation (5, 10, 15 μg sequentially) on days 0, 28 and 56 (Figure 1C). We also included a third group of mice which received an adjuvanted recombinant *Pf*MSP2 vaccine [17] as a reference for the mRNA *Pf*MSP2 vaccine performance and dose optimisation following the same regimens and sampling schedule. Sera were collected at each vaccine dose and two weeks after the final dose on day 70 (peak vaccine response), and were assessed for vaccine-induced antibodies by enzyme-linked immunosorbent assay (ELISA). Vaccine-induced IgG to *Pf*MSP2 3D7 increased after each dose in all vaccine groups (Figure 1D). Interestingly, there was no significant difference between the recombinant and the two mRNA *Pf*MSP2 regimens indicating higher mRNA doses were not required to achieve optimal IgG induction. Next, the sera from the individual mouse groups were pooled and used to detect recombinant *Pf*MSP2 3D7 in western blots, and the groups showed comparable detection (Figure 1E). To establish that vaccine-induced antibodies recognise native *Pf*MSP2 on the merozoite surface, whole intact merozoites were isolated from *P. falciparum* blood-stage culture and used in ELISA. Both recombinant and mRNA *Pf*MSP2 vaccines induced IgG at peak (day 70) that binds native *Pf*MSP2 on the merozoite surface without a significant difference [28], indicating recognition of key native epitopes (Figure 1F).

**Figure 1.**
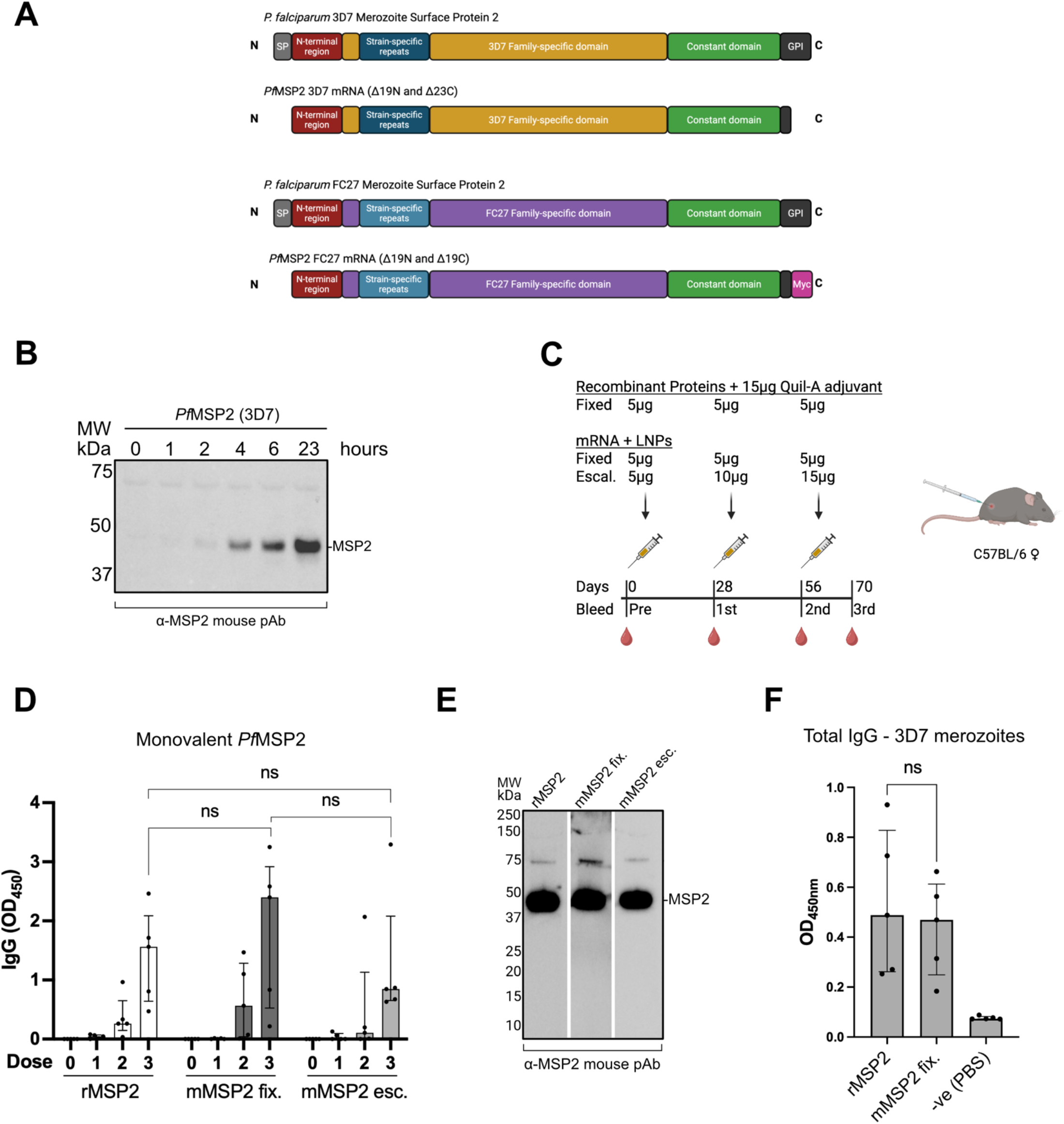
*Pf*MSP2 3D7 mRNA vaccine evaluation. (A) Schematics representing the full-length *Pf*MSP2 3D7 and FC27 alleles, and the truncated mRNA constructs with the deletion of the N-terminal signal peptide (SP) and C-terminal GPI anchor motif. *Pf*MSP2 FC27 mRNA construct contained a Myc tag for mammalian expression detection in WB. (B) Expi293F cells were transfected with naked *Pf*MSP2 3D7 mRNA and cell lysate was collected at multiple time-point post-transfection. The samples were processed under reducing conditions for western blot, and protein expression was detected using mouse α-*Pf*MSP2 polyclonal IgG (1/1000 dilution). (C) Schematic of mouse immunisation regimens and timeline for blood collection (recombinant *Pf*MSP2: rMSP2; mRNA *Pf*MSP2 fixed dose: mMSP2 fix; mRNA *Pf*MSP2 dose escalation: mMSP2 esc). (D) Adjuvanted recombinant and mRNA-LNP *Pf*MSP2 immunogenicity ELISA analysis of IgG in mouse sera at 1/25600 dilution (n=5 per group). Error bars represent median ± IQR and assays performed in duplicates. Statistical analysis was performed using Kruskal-Wallis test; ns is not significant (p>0.05). (E) Western blot detection of 250 ng of recombinant *Pf*MSP2 loaded into each lane and detected by pool mouse sera polyclonal IgG from immunisation regimens (1/1000 dilution). (F) ELISA analysis of mouse sera IgG from day 70 at 1/100 dilution (n=5 per group) detecting native *Pf*MSP2 on whole merozoites. Error bars represent median ± IQR and assays performed in duplicates. Statistical analysis was performed using Mann-Whitney test; ns is not significant (p>0.05).

### PfMSP2 mRNA vaccines induce IgG with Fc-mediated functional activity

The profiles of murine IgG subclasses induced by recombinant and mRNA *Pf*MSP2 vaccines were examined to understand the functional or protective potential of vaccine responses [29]. The cytophilic IgG subclasses in C57BL/6 mice with human analogues are IgG1 (huIgG4), IgG2b (huIgG3), IgG2c (huIgG1), and IgG3 (huIgG2). All murine IgG subclasses confer protective functions, however IgG1 exhibits the lowest functional activity relative its high abundance in serum [22, 30, 31]. Testing IgG at the peak response point (day 70), the three vaccine regimens showed comparable subclass profiles with IgG1 at higher magnitude, and moderate IgG2b/c induction, but low IgG3 levels (Figure 2A). In studies of human immunity, it has been shown that antibodies to *Pf*MSP2 have Fc-mediated functions including complement fixation and activation, which can lyse merozoites and inhibit erythrocyte invasion, antibody-dependant cellular inhibition (ADCI) and opsonic phagocytosis of merozoites, mediated by IgG interactions with Fcγ receptors [9–11, 17, 29, 32, 33]. Therefore, we assessed similar Fc-mediated functions of vaccine-induced mouse antibodies. There were multiple functional antibody activities induced by vaccination among all mouse groups. Notably, the murine IgG showed strong binding to human Fcγ receptors (I and IIa), which are activating receptors promoting phagocytosis and other functions. We did not assess FcγRIII binding because mouse IgG has weak affinity to human FcγRIII [34]. The mouse antibodies also demonstrated strong fixation of human C1q, which is the first essential step in classical complement activation. All functional activities were broadly comparable between groups, with mRNA *Pf*MSP2 fixed-dose sera tending to have the highest activity, although this was not significant due to biological variation (Figure 2B). Additionally, a pool of serum from mice vaccinated with mRNA *Pf*MSP2 was used to opsonise *Pf*MSP2 3D7 coated fluorescent beads and examine IgG-mediated phagocytosis *in vitro*. The monocytic THP-1 cells showed substantial phagocytosis of opsonised beads relative to the negative control (PBS only) indicating Fc receptor recognition and function (Figure 2C). This supports the cross-reactivity of murine IgG and human Fc receptors, with strong correlations between the *in vitro* binding to recombinant human Fcγ receptors observed of vaccine-induced antibodies, most significantly functional murine IgG2c (Figure 2D).

**Figure 2.**
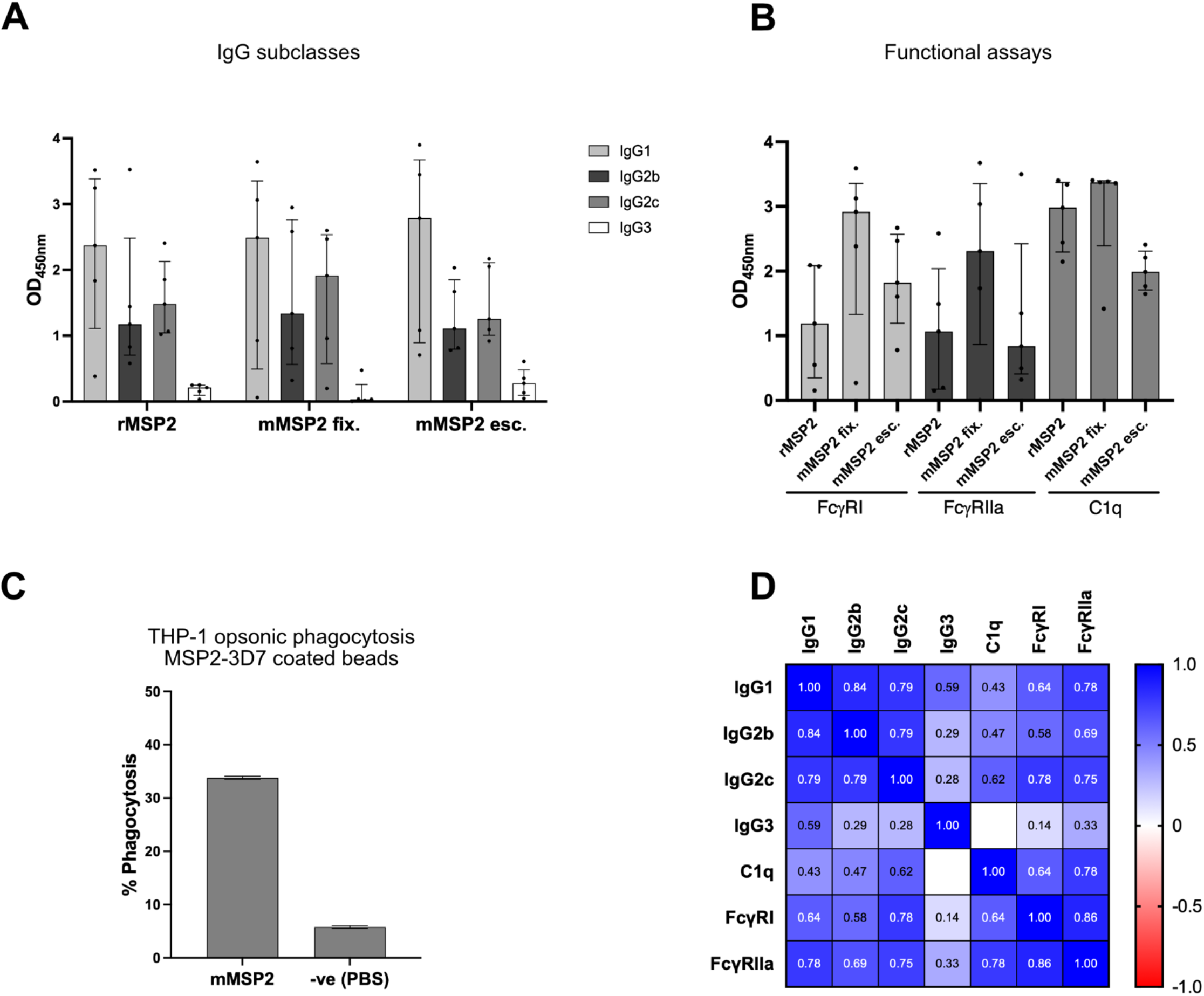
Functional activity of *Pf*MSP2 3D7 monovalent mRNA vaccine IgG. (A) ELISA analysis of mice IgG subclasses in day 70 sera (n=5 per group). Assays were performed in duplicates (1/6400 dilution) and error bars represent median ± IQR. (B) Functional assays of murine IgG from day 70 sera for human FcγRI binding (1/12800 dilution), human FcγRIIa binding (1/400 dilution), and human C1q engagement (1/100 dilution). Assays were performed in duplicates and error bars represent median ± IQR; two-way ANOVA statistical test showed no significant differences between regimens for all functional assays (p>0.05). (C) Phagocytosis of *Pf*MSP2-coated fluorescent beads by THP-1 monocytes using mMSP2 pooled sera samples (n=5 mice) and 1/60 dilution with PBS for negative controls. Assays performed in duplicates and error bars represent median + 95% CI. (D) Correlation matrix (values are Spearman’s rho) of mouse IgG subclasses at peak and function (day 70; n=15 mice) with +1 indicating strong positive correlation, 0 indicating no correlation, and -1 indicating strong negative correlation. All correlations were p<0.05 except IgG3.

### Bivalent PfMSP2 mRNA vaccine incorporating the two major alleles elicits high functional IgG responses

In *P. falciparum* populations, *Pf*MSP2 occurs as two major allelic types and antibodies to *Pf*MSP2 typically have a degree of allele-specific reactivity. Therefore, an important property for a protective *Pf*MSP2 vaccine is the ability to generate an immune response to both major alleles (3D7 and FC27) [17, 33]. To achieve this, the *Pf*MSP2 FC27 allele was codon-optimised and truncated (Δ19N and Δ19C) to remove the N-terminal leader sequence and GPI anchor respectively, with the addition of a C-terminal Myc tag (Figure 1A). Following expression validation of the *Pf*MSP2 FC27 allele (Figure S1), the two alleles were encoded by mRNA and formulated as mRNA-LNPs individually then mixed in a 1:1 ratio and used to immunise mice (n=5). The regimen in this study included 3 × 10 μg doses (5 μg per allele) and sera collection at regular intervals for analysis (Figure 3A). The vaccine induced high and equivalent IgG responses against both alleles, with vaccine-induced IgG highest titre after the 3^rd^ dose (Figure 3B). Additionally, there was a strong induction of the functional murine subclasses in the bivalent mRNA *Pf*MSP2 vaccine and high functional activity by binding to human Fcγ receptors and human C1q (Figure 3C-D). This activity was measured using recombinant *Pf*MSP2 3D7 and FC27, and these antibodies are expected to bind native *Pf*MSP2 on the parasite surface as observed in the monovalent mRNA vaccine. These results suggest the vaccine could generate potent and strain-transcending *Pf*MSP2 antibodies capable of protective immunity [12, 22].

**Figure 3.**
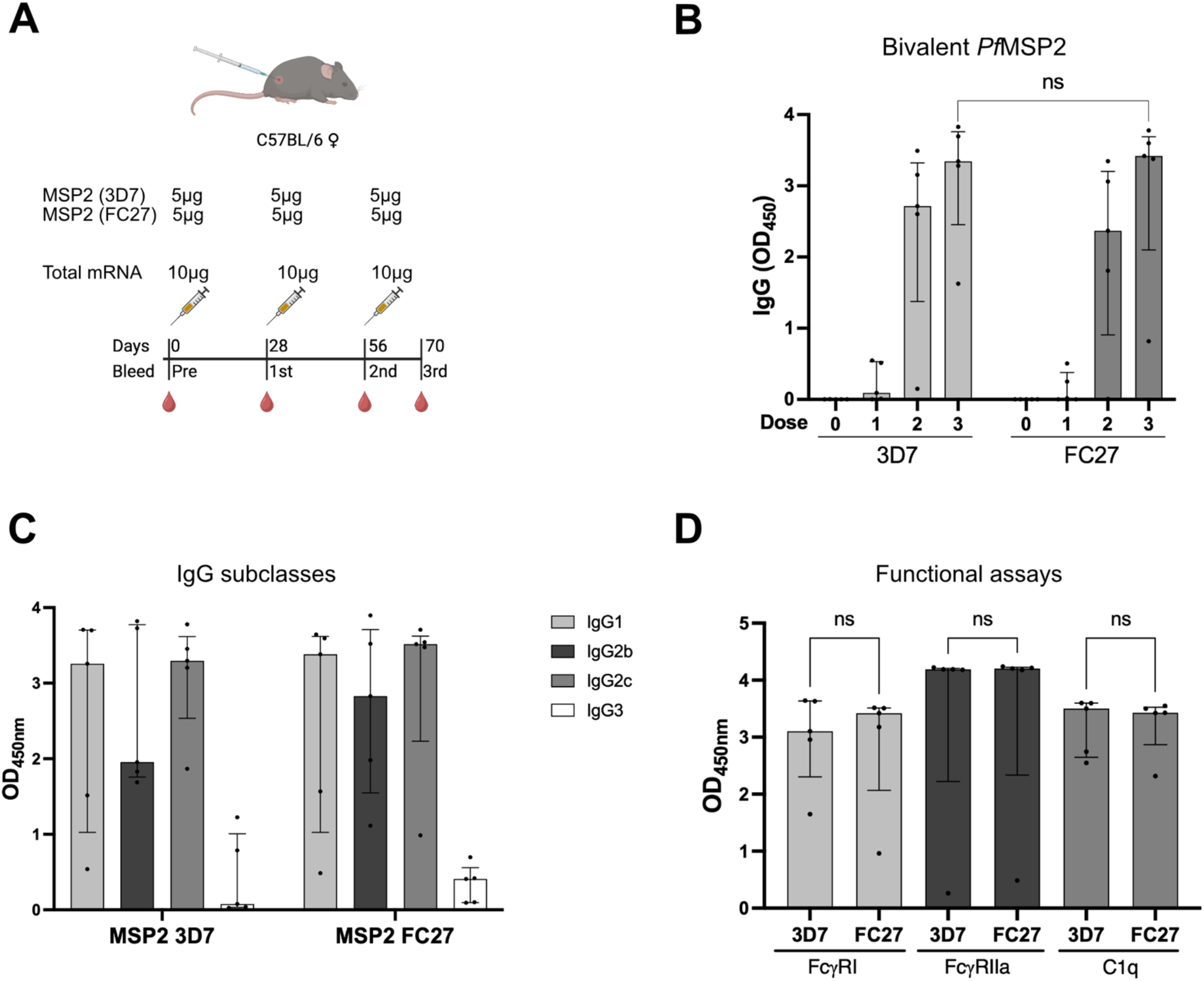
*Pf*MSP2 bivalent (3D7 and FC27) mRNA vaccine evaluation. (A) Schematic of mice immunisation regimens and timeline of blood collection for bivalent *Pf*MSP2 formulations (3D7 and FC27 alleles). (B) Bivalent mRNA-LNP (3D7 and FC27) immunogenicity ELISA analysis of IgG in total mouse sera at 1/6400 dilution (n=5). Error bars represent median ± IQR and assays performed in duplicates. Statistical analysis was performed using Mann Whitney test; ns is not significant (p>0.05). (C) ELISA analysis of murine IgG subclasses in sera at day 70 (n=5). Assays were performed in duplicates (1/3200 dilution) and error bars represent median ± IQR. (D) Functional assays of mice IgG from day 70 sera for human FcγRI binding (1/12800 dilution), human FcγRIIa binding (1/400 dilution), and human C1q engagement (1/800 dilution). Assays were performed in duplicates and error bars represent median ± IQR. Statistical analysis was performed using Kruskal-Wallis test; ns is not significant (p>0.05).

## Discussion

In this study we demonstrate a strategy to target the merozoite surface using mRNA vaccines to generate antibodies with Fc-mediated functional activity. We designed mRNA vaccines based on *Pf*MSP2 because it is an abundant merozoite surface antigen and a known target of antibodies that promote complement activation to inhibit blood-stage replication and mediate opsonic phagocytosis of merozoites [9–12]. Furthermore, antibodies to *Pf*MSP2 have been associated with protection in studies of naturally-acquired and vaccine-induced immunity [10, 12]. We formulated monovalent and bivalent vaccines including the two major allelic isoforms of *Pf*MSP2 and found the vaccine effectively generated antibodies with Fc-mediated functional activities to both alleles in preclinical studies.

Validating the expression of *Pf*MSP2 in mammalian cells was essential for progressing the mRNA vaccine into preclinical animal studies. The mRNA encoding *Pf*MSP2 3D7 and FC27 was codon-optimised for mammalian translation, and expression of both antigens was detected indicating compatibility of *Pf*MSP2 with the mRNA platform. The mRNA-LNP formulation showed immunogenicity for both monovalent (3D7) and bivalent (3D7 and FC27) vaccines in mice, comparable to the adjuvanted recombinant antigen vaccine. The efficiency of mRNA vaccine translation *in vivo* has been reported to be 1-2% due to endosomal escape limitations, which equates to an estimated ≥100 ng of *Pf*MSP2 for the 5 μg mRNA-LNP vaccine dose compared to the 5 μg of *Pf*MSP2 when given as a recombinant protein formulated with adjuvant [35]. Remarkably, the immunogenicity of the mRNA vaccine was comparable to the recombinant protein vaccine, and there was no improved immunogenicity using higher doses mRNA indicating dose-sparing advantages. However, further dose optimisation studies and larger sample sizes would be needed to more formally evaluate dosing, compare the mRNA and recombinant protein platforms, and examine biological variability within and between regimens.

The mRNA vaccine-induced antibodies by monovalent and bivalent formulations showed strong functional activity, including complement fixation, interactions with FcγRI and FcγRIIa, and opsonic phagocytosis [34, 36]. This functional antibody profile is similar to what has been reported for antibodies in naturally-acquired immunity and clinical trials of a recombinant *Pf*MSP2 protein vaccine in humans [11, 12]. These Fcγ receptor engagement and complement activation functions are key to parasite inhibition and clearance *in vitro* and *in vivo* [9, 11, 12, 37]. The IgG2c and IgG2b subclasses of C57BL/6 mice, which have Fc-mediated functional activities, were collectively higher than murine IgG1, which can conversely confer inhibitory activity of downstream immune functions [38]. Thus, it was promising that the bivalent formulation showed the same induction of antibodies to both allelic forms.

Both cross-reactive and allele-specific antibodies to *Pf*MSP2 have been observed in naturally-acquired and vaccine-induced immunity in humans and cross-reactivity may be partly attributed to antibodies against the conserved C-terminal region [12]. Naturally-acquired antibodies were reported to more prominently target the variable allele-specific regions of *Pf*MSP2 than vaccine-induced antibodies [12, 33]. Further investigation of responses induced by the bivalent vaccine to map key epitopes targeted by functional and cross-reactive antibodies may be valuable to fine-tune *Pf*MSP2 construct design. It is also important to understand the kinetics of cellular immune responses that may provide insights into vaccine longevity and potential protection. Evaluating the protective efficacy of *Pf*MSP2-based vaccines is not possible with current mouse models of malaria because *Pf*MSP2 or an orthologue is not present in rodent malaria species such as *Plasmodium berghei*, and *P. falciparum* does not infect rodents. Furthermore, the very low activity of complement fixation in laboratory mice and different use of *in vitro* Fc-receptors and immune cell strategies compared to humans pose further challenges [4]. Therefore, evaluating *Pf*MPS2 vaccines will require non-human primate studies or controlled human malaria infection (CHMI) vaccine trials. 225

## Conclusions

In our study we showed that the mRNA-LNP platform can be used for monovalent and bivalent formulations that induce functional IgG against *Pf*MSP2, further supporting *Pf*MSP2 as a vaccine candidate and the broader concept of targeting merozoite surface proteins in malaria vaccine strategies. We showed that a 3-dose regimen induced peak immunity comparable to the adjuvanted recombinant protein. Another lead merozoite vaccine based on *Pf*Rh5 induces growth-inhibitory antibodies and shows modest efficacy in malaria endemic populations. However, studies of naturally-acquired immunity point to the importance of antibody Fc-mediated functions in protective immunity. Therefore, combining *Pf*Rh5 with *Pf*MSP2, or other antigen combinations with *Pf*MSP2, may be an effective strategy to induce antibodies with both growth-inhibitory and Fc-mediated functional activities for greater protective efficacy. Further, a highly effective blood-stage vaccine combined with the leading sporozoite antigen, *Pf*CSP, could reduce the risk of initial infection and subsequent disease to achieve high-level protection against malaria.

## Materials and Methods

### Production of naked mRNA and mRNA-LNP vaccines

Codon-optimised *P. falciparum* MSP2 3D7 and FC27 sequences (GenBank accession number JN248383 and JN248384 without the C-terminal poly-Histidine tag) were synthesised as described previously [25]. The mRNA vaccines in this study used Moderna’s LNP formulation (SM102) in 8.8% (w/v) sucrose in 25 mM Tris buffer (pH 7.4). All mRNA synthesis, LNP formulation, and characterisation were performed by mRNA Core at the Monash Institute of Pharmaceutical Sciences, Australia.

### Expression and validation of naked mRNA

The expression of naked mRNA was performed in Expi293F suspension cells (Gibco) using TransIT-mRNA (Mirus) transfection kit. Briefly, 5 μg of naked mRNA was added to 683 μL of Opti-MEM GlutaMAX (Gibco), then 7 μL of the Booster reagent was added and gently mixed, and finally 7 μL of the TransIT reagent was added and mixed. The transfection mix was incubated for 2 minutes at RT then added dropwise to 5 mL of Expi293F cells in a 6-well plate at density 1 × 10^6^ cell/mL. The cells were incubated shaking in high humidity at 37°C and 8% CO_2_, and 1 mL samples were collected at multiple time-points for analysis via NuPAGE and western blot.

### PfMSP2 production

Recombinant *Pf*MSP2 3D7 and FC27 were manufactured at GroPep Pty Ltd (Adelaide, Australia) by expression in *E. coli* and purified from lysate by chelating, anion-exchange, and reverse-phase chromatography. The purified protein was stored at -80°C in 10 mM acetic acid to resolve amyloid-like fibrils formation [17].

### Study design and mice immunisation

The evaluation of the *Pf*MSP2 mRNA-LNP and adjuvanted recombinant *Pf*MSP2 vaccines was conducted using female C57BL/6 mice aged 8 – 12 weeks (n=5 per group). The mice were immunised with two regimens for mRNA-LNP, fixed dose (3 × 5 μg; fix. group) and dose-escalation (5 μg, 10 μg, 15 μg; esc. group). The recombinant protein vaccine followed a single regimen (3 × 5 μg *Pf*MSP2 protein and 15 μg Quil-A adjuvant; Croda). The mice were immunised at 28 days intervals with prior blood collection proceeding the injection. The immunisation and blood collection schedules are shown in Figures 1B and 3A. Mice received all immunisations via intramuscular injections to the calf muscle. The first two collections are retro-orbital bleeds, the subsequent collections are mandible bleeds, and the terminal collection is a cardiac bleed (following inhaled anaesthetic euthanasia). All mice were housed, cared for, and handled (injection and bleeding) by the WEHI Antibody Facility in Bundoora.

### Culture and isolation of P. falciparum merozoites

Parasite culture (*Plasmodium falciparum* 3D7) and merozoite isolation were performed as previously described [9, 39].

### Culture of THP-1 monocytes and phagocytosis assay

These methods were previously described and optimised [10, 12, 40, 41]. Briefly, THP-1 cells were counted and resuspended to a density of 5 × 10^6^ cell/mL in THP-1 complete medium. Recombinant *Pf*MSP2 3D7 was coupled with amine-modified fluorescent latex beads (Sigma-Aldrich) and used at 0.75 × 10^6^ beads/mL. For opsonisation, pooled sera from 5 mice were diluted to a 50 µL final volume using 1% (w/v) Bovine Serum Albumin (BSA; Bovogen) in 1 × PBS. The diluted sera were then incubated with the coated beads for 1 hour at RT. Following washing of beads, THP-1 cells were added and incubated at 37°C and 8% CO_2_ for 40 minutes in the dark. Following washing and fixation, the cells were resuspended in 1% (w/v) BSA in 1 × PBS. Finally, quantification of phagocytosis was determined using flow cytometry in 96-well format on BD FACS Canto II (BD Biosiences). PBS only was used in negative controls.

### NuPAGE expression analysis

To assess the expression of *Pf*MSP2 mRNA, the transfection samples were run on a gel under reducing conditions. Briefly, whole cell samples were prepared to contain 100 mM dithiothreitol (Thermo Fisher Scientific), and 1 × NuPAGE LDS Sample Buffer (Invitrogen). Samples were incubated at 95°C for 5 minutes, vortexed for 30 seconds, then loaded onto a 4-12% Bis-Tris gel (Invitrogen) with Precision Plus Protein Dual Color Standards (Bio-Rad) and ran at 135 V and 200 mA for 1.5 hours in 1 × MES running buffer (Invitrogen). The gel was then analysed by western blot.

### Western blot analysis

The polyacrylamide gel was transferred onto a nitrocellulose (NC) membrane using iBlot 2 Transfer Stacks (Thermo Fisher Scientific) at a current of 20V for 7 minutes (default setting P3). The membrane was blocked with 1% (w/v) BSA in 0.05% (v/v) Tween20 (Sigma-Aldrich) in 1 × PBS (PBS-T) overnight at 4 °C. The membrane was probed with mouse anti-*Pf*MSP2 3D7 primary antibodies (1/1000), and goat anti-mouse IgG horseradish peroxidase (Merck Millipore) secondary antibody (1/2000) in 1-2% (w/v) BSA at RT with extensive washing with PBS-T between steps. The SuperSignal West Pico PLUS Chemiluminescent Substrate (Thermo Fisher Scientific) was used for detection. The membrane was then visualised using the ChemiDoc MP (Bio-Rad).

### ELISA and plate-based assays

ELISA protocols were performed as described previously for IgG, C1q fixation, and human Fc receptor binding [10, 12, 29, 37, 42, 43]. All plate washing was performed between each step with 0.05% (v/v) Tween in 1 × PBS using the automated 405 LS microplate washer (Agilent Biotek). All assays were normalised across plates for individual experiments.

To measure IgG, 0.5 µg/mL of recombinant *Pf*MSP2 3D7/FC27 was used to coat flat-bottom 96-well Nunc Maxisorp (Thermo Fisher Scientific) microplates overnight at 4°C. The plates were then blocked with 0.1% (w/v) casein (Sigma-Aldrich) in 1 × PBS and incubated at 37°C for 2 hours. Diluted mouse serum samples in 0.1% (w/v) casein in 1 × PBS were then added and incubated for 2 hours at RT, followed by HRP-conjugated antibodies (goat anti-mouse IgG HRP, Merck Millipore; anti-mouse IgG subclasses HRP, Southern BioTech) diluted in 0.1% (w/v) casein in 1 × PBS and incubated for 1 hour at RT. Finally, the plates were incubated with TMB liquid substrate (Life Technologies) and incubated in the dark for up to 1 hour at RT while the colour-change developed. The reaction was stopped using 1% (v/v) H_2_SO_4_ in distilled H_2_O and OD values were immediately read at 450nm using a Multiskan GO plate reader (Thermo Fisher Scientific).

To measure vaccine-induced IgG in mouse sera binding to merozoites, the standard IgG ELISA protocol was followed with the following modifications: coating wells with 5 × 10^4^ merozoites/mL diluted in 5% (w/v) skim milk in PBS-T at 37°C for 2 hours. All dilution steps of serum and antibodies were performed using 5% (w/v) skim milk in PBS-T, and plate blocking using 10% (w/v) skim milk in PBS-T.

To measure C1q fixation, modified ELISA protocols were performed as described previously. The plates were coated with 1 µg/mL of recombinant antigen and sera was incubated following the previous steps. Human C1q (Merck Millipore) was diluted to 10 µg/mL with 0.1% (w/v) casein in 1 × PBS and incubated for 30 minutes at RT, then rabbit anti-C1q IgG (Burnet Institute) [44] diluted with 0.1% (w/v) casein in 1 × PBS and incubated for 1 hour at RT. The goat anti-rabbit IgG HRP (Bio-Rad) was diluted with 0.1% (w/v) casein in 1 × PBS and incubated for a further 1 hour at RT before the substrate was added, reaction stopped, and absorbance measured.

To measure IgG binding to human FcγRI and FcγRIIa, the same standard IgG ELISA steps were followed with the exception of using 1% (w/v) BSA in 1 × PBS for blocking, and sample and reagent dilution. Following sera incubation, 0.125 µg/mL of biotinylated FcγRI 10256-H27H-B (Sino Biological) or 0.2 µg/mL dimeric FcγRIIa (Burnet Institute) [45] was added and incubated at 37°C for 1 hour. For detection of FcγRs, Streptavidin-HRP (ThermoFisher Scientific) diluted 1/10000 was added and incubated at 37°C for an additional hour, followed by substrate reaction and absorbance measurement.

### Statistical analysis

All statistical analyses and graph generation for this study were conducted using GraphPad Prism 11 (MacOS) software. Mann-Whitney test was used to compare two groups, while the Kruskal-Wallis test was used for multiple groups (more than two) comparisons. A two-way ANOVA test with mixed effects was applied to account for the dependence of data points over time. Spearman’s rho was calculated to assess correlations between parameters.

## Acknowledgements

We thank Bruce Wines for providing Expi293F-BirA cells and the FcγRIIa H131clone. We also thank Paul Masendycz, Ridouan Bouhbouh, and Myha Huynh at WEHI Antibody Facility for their support with our animal studies. We thank Robin Anders for his valuable advice on working with recombinant and native *Pf*MSP2. Schematics were created using BioRender. Burnet Institute is located on the traditional lands of the Boonwurrung people of the Kulin nation.

## Ethics

Burnet Institute malaria antigens mRNA handling approval (IBC 40702). WEHI Antibody Facility animal and mRNA handling in Bundoora; Scientific Procedure Premises Licence (SPPL201904), and ethics approval (2020.019 and 2023.012).

## Author contribution

AT and JB designed the study. AT, TR, TH, and SC conducted the experiments. AT, TR, and TH were involved in data analysis with guidance from LR, LK, SC, CD, and JB. SF, CLL, RSM, and CP synthesised and formulated mRNA and mRNA-LNP. AT led writing the manuscript with input from other authors. All authors reviewed and approved the final version.

