## Supplemental Figure 1 for "Targeting a malaria merozoite surface protein with mRNA vaccine generates multifunctional antibodies"

Thomas A. et al, 2026

**Supplementary material**

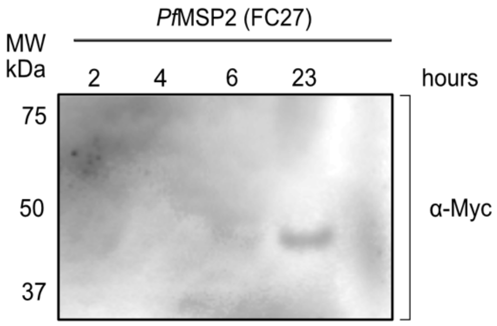


**Figure S1. *Pf*MSP2 FC27 mRNA expression validation**

Western blot of time-course total Expi293F cell lysate expressing *Pf*MSP2 FC27. The NC membrane was probed with mouse $\alpha$-Myc monoclonal antibody (Cell Signaling; 1/1000 dilution).
